# Differential taphonomic effects of petroleum seeps and karstic sinkholes on ancient dire wolf teeth: Hydrocarbon impregnation preserves fossils for chemical and histological analysis

**DOI:** 10.1101/2021.01.04.425345

**Authors:** Sabrina B. Sholts, Leslea J. Hlusko, Joshua P. Carlson, Sebastian K. T. S. Wärmländer

## Abstract

Histological analysis of teeth can yield information on an organism’s growth and development, facilitating investigations of diet, health, environment, and long-term responses to selective pressures. In the Americas, an extraordinary abundance of Late Pleistocene fossils including teeth has been preserved in petroleum seeps, constituting a major source of information about biotic changes and adaptations at the end of the last glacial period. However, the usefulness of these fossils for histological studies is unclear, due to the unknown taphonomic effects of long-term deposition in petroleum. Here, we compare histological and chemical analyses on dire wolf (*Canis dirus*) teeth obtained from two different environments, i.e. a petroleum seep (Rancho La Brea tar pits, California) and a carstic sinkhole (Cutler Hammock sinkhole, Florida). Optical and scanning electron microscopy (SEM) together with X-ray diffraction (XRD) analysis revealed excellent preservation of dental microstructure in the seep sample, and the petroleum-induced discoloration was found not to interfere with the histological and chemical examination. By comparison, teeth from the sinkhole sample showed severe degradation and contamination of the dentine by exogenous substances. These results indicate that petroleum seep assemblages are useful, or even ideal, environments for preserving the integrity of fossil material for chemical and histological analysis.

## 1. INTRODUCTION

The hard tissues of teeth are remarkably tough, and unlike bone, are not all continually replaced during life. Teeth can therefore preserve a rich temporal record of growth and development, and the effects of diet, health, and the environment on these patterns (Antoine et al., 2009; Dean, 2006). For extinct organisms in particular, whose biological remains are often scarce, teeth may be the best available source of information for studies of ontogeny, life history, evolution, and/or taxonomy (Smith, 2008). However, certain features of tooth formation are only present in the internal microstructure of dental tissues. Analyzing these features typically requires destructive histological methods, such as cutting the tooth into cross-sections. Ideally these destructive methods should only be used on teeth with well-preserved microstructure (Hillson, 2005), as the histological results may be compromised by external factors such as taphonomic degradation and result in the unnecessary loss of a limited resource.

Abundant remains of extinct organisms have been preserved as fossils in petroleum seeps, such as the Rancho La Brea “tar pits” in California. These seeps form when natural hydrocarbons are pressured up from their underground reservoirs to form viscous petroleum sediments on the earth’s surface. When animals or their remains become entrapped in the sticky asphalt, the resulting accumulation of bones turns these seeps into fossil deposits. In the Americas, petroleum seeps have yielded large quantities of terrestrial vertebrate faunal remains from the Late Pleistocene, a period of significant climatic change and megafaunal extinctions (Haynes, 2008). Among these fossils are the earliest evidence of the grey fox (*Urocyon cinereoargenteus*) in South America (Prevosti and Rincón, 2007), the oldest fossil mammals in Cuba (Jull et al., 2004), and one of the few Paleoamerican human skeletons found in North America (Berger, 1975).

Vertebrate fossils from petroleum seeps are recognizable by the infiltration of hydrocarbons into the hard tissue matrices of bones and teeth. Although heavy asphaltic surface coating can be manually removed or dissolved in various (toxic) solvents (Shaw, 1982), particles causing dark discoloration permeate the entire specimen. Petroleum impregnation does not appear to chemically react with the organic or apatite structures of the bone tissues (Shelton, 1994), and seems to act as an ideal preservative for bone collagen (Ho et al., 1969). Contaminating petroleum can be removed from bone material without significant impact on either nuclear or mitochondrial DNA (Janczewski et al., 1992), and separated from amino acids to facilitate radiocarbon dating (Ho et al., 1969; Jull et al., 2004). However, early studies indicate that discoloration from petroleum impregnation has varying effects on histology, ranging from negligible (Cook et al., 1962; Wyckoff and Doberenz, 1965) to obstructive (Ørvig, 1978).

Here, we evaluate the suitability of fossil teeth from petroleum seeps for dental histology using light microscopy and scanning electron microscopy (SEM) with energy-dispersive x-ray spectroscopy (EDS). Contemporaneous and conspecific teeth from a non-seep deposit are used to compare preservation and visibility of various microscopic features and tissue properties.

## 2. TOOTH DEVELOPMENT AND STRUCTURE

Teeth consist of three calcified tissues of different origins, function, and chemical composition, i.e. enamel, dentine, and cement, situated in an organic collagen matrix. The hardness of these tissues is due to a predominant inorganic component of calcium phosphate minerals, mostly in the form of long and narrow apatite crystallites. Mature enamel is almost entirely inorganic, with densely packed and highly organized crystallites that are significantly longer than those of bone and dentine (Hillson, 2005:155). As the hardest tissue in the mammalian body, enamel protects the tooth from the oral environment by covering the entire crown. Dentine constitutes the majority of the tooth, bounded by enamel and cement at its outer border and the pulp cavity at its interior border. Dentine crystallites are seeded in the collagen matrix, which is secreted in mats of fine fibers (Hillson, 2005:184). Due to its higher organic content and structure, dentine is more porous and less tough than enamel. On the dentine and enamel surfaces, cement is formed as a softer, bone-like substance of variable thickness and distribution (Hillson, 2005:193). Its primary function is to provide a tissue attachment for the periodontal ligament, which anchors the tooth to the bony walls of the socket and thus holds its position in the mandible or maxilla.

Dental tissues are formed by accretional processes that begin before birth and continue through adolescence, preserving incremental markings of an individual’s developmental history (Dean, 1987, 2006; Hillson, 2005; Smith, 1989, 1991; Smith, 2008). Although the slowly forming cement provides a record of widely spaced increments of seasonal or annual growth, the more quickly forming enamel and dentine reflect growth patterns of much shorter periodicity (Dean, 1987). Daily (circadian) variation in the secretory activity of enamel-forming cells (ameloblasts) produces fine transverse striations that occur along the length of enamel prisms. Slight accentuations of these cross-striations are known as brown striae of Retzius, and represent a nearly weekly (circaseptan) rhythmic disturbances in crown formation (Dean, 1987). Cross-striations and Retzius lines also appear to correspond to short-period and long-period features in the dentine, known as von Ebner’s lines and Andersen lines respectively (Dean and Scandrett, 1996).

In addition to incremental markings, enamel microstructure displays a number of aperiodic features that are visually distinctive and functionally relevant. The apatite crystallites in enamel have many different orientations, which limit the extension of any potential fracture (Boyde, 1964:170). This poly-orientation can be observed in the change in orientation of crystals across individual prisms and within inter-prismatic regions, or in the decussation of prisms in Hunter-Schreger bands (Boyde, 1964:170). Prism patterns have been differentiated on the basis of cross-sectional outlines of prism shape and organization in mature mammalian enamel (Boyde, 1964), and associated with different enamel deposition rates and taxonomic groups (Dumont, 1995).

## 3. PETROLEUM CHEMISTRY AND PROPERTIES

Petroleum (crude oil) consists of hydrocarbons mainly in cyclic configurations, known as “aromats” due to the strong and pungent aroma of these molecules (Fig. 1). Typical properties of these polycyclic aromatic hydrocarbons (PAHs) include low volatility at room temperature, poor solubility in water, and high lipophilicity. These characteristics become more pronounced with increasing molecular weight (i.e., greater number of rings), which is accompanied by increased melting/boiling points, decreased resistance to oxidation and reduction, and decreased vapor pressure (Eisler, 2000).

**Figure 1.**
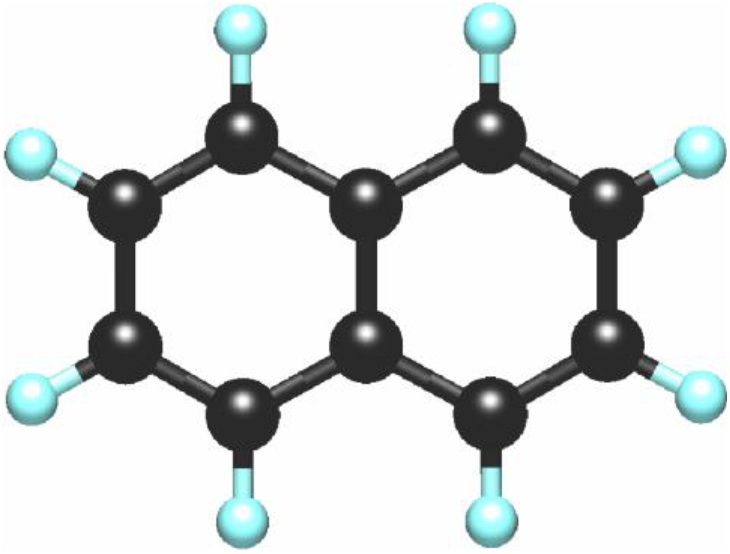
Ball and stick model of the three-ringed planar polycyclic aromatic hydrocarbon (PAH) *phenanthrene*, one of the most common hydrocarbons in petroleum from the Rancho La Brea (RLB) tar pits.

The hydrocarbons in petroleum are formed by breakdown of dead organisms under certain conditions in the earth’s crust. The organic material first undergoes anaerobic enzymatic degradation and microbiological decomposition in sediment (Albrecht and Ourisson, 1971). The organic sediment then sinks, becomes buried in a basin, and is subjected to gradual rises in temperature and pressure (Albrecht and Ourisson, 1971). When heated to the right temperatures, the insoluble fraction of sedimentary organic matter, known as kerogen, is converted into polycyclic hydrocarbons (Vandenbroucke and Largeau, 2007). Petroleum in liquid or semi-solid states is also known as bitumen or asphalt, and should not be confused with tar, which is man-made and produced by destructive distillation of e.g. coal, wood, petroleum, or peat. However, due to the similar black and semi-solid appearance of petroleum and tar, petroleum seeps are often referred to as “tar seeps” or “tar pits”.

The chemistry of petroleum is important for understanding its taphonomic effects on bones and teeth. Even though many hydrocarbons are harmful to humans and animals (Sholts et al., 2017; Wallin et al., 2017; Westerholm et al., 2001; Wärmländer et al., 2011), they rarely interact chemically with other molecules (such as collagen and apatite) due to their low reactivity (Ho et al., 1969; Shelton, 1994), and they do not degrade quickly due to their high stability. Encased in the anoxic environment of a petroleum seep, organic matter may therefore be protected against some diagenetic effects. Furthermore, petroleum itself can be an interesting subject-matter for anthropological and paleontological investigations, as it e.g. can help provenance asphaltic fossils. Because of varying geological conditions, all seeps have their own signature ratios of different PAHs (Wärmländer et al., 2011) and also slightly different isotope and trace element compositions, allowing petroleum from a specific seep (e.g., Rancho La Brea) to be uniquely identified (Brown et al., 2014; Wendt and Lu, 2006).

## 4. MATERIALS

The total sample for this study consisted of 25 teeth of 19 dire wolves (*Canis dirus*) (Table 1). From this sample, 15 teeth from 11 individuals were selected for destructive analysis, and the remaining ten teeth were left untreated for non-destructive examination. Teeth were obtained from two museum collections of Rancholabrean age faunal remains, i.e. finds from the Rancho La Brea (RLB) tar pits and from the Cutler Hammock (CH) sinkhole. In addition, teeth from a modern gray wolf (Canis lupus) were investigated as comparative material.

**Table 1.** Total sample of dire wolf teeth used for this study, from the University of California Museum of Paleontology (UCMP) and the Florida Museum of Natural History (FLMNH).

| Museum | ID | Site | Tooth type | Sectioned |
| --- | --- | --- | --- | --- |
| FLMNH | 135885 | Cutler Hammock | M1 | no |
| FLMNH | 276780 | Cutler Hammock | C | yes |
| FLMNH | 276782 | Cutler Hammock | C | no |
| FLMNH | 276784 | Cutler Hammock | M1 | no |
| FLMNH | 276785 | Cutler Hammock | M1 | no |
| FLMNH | 276786 | Cutler Hammock | P4 | no |
| FLMNH | 276788 | Cutler Hammock | P2 | yes |
| FLMNH | 276791 | Cutler Hammock | M1 | no |
| FLMNH | 276797 | Cutler Hammock | P2 | yes |
| FLMNH | 276798 | Cutler Hammock | M2 | no |
| UCMP | 215833 | Rancho La Brea | P2 | yes |
| UCMP | 215834 | Rancho La Brea | P2 | yes |
| UCMP | 215837 | Rancho La Brea | P1 | yes |
| UCMP | 215840 | Rancho La Brea | P2 | yes |
| UCMP | 215842 | Rancho La Brea | C | yes |
| UCMP | 215842 | Rancho La Brea | P3 | no |
| UCMP | 215842 | Rancho La Brea | P4 | yes |
| UCMP | 215842 | Rancho La Brea | P4 | no |
| UCMP | 215843 | Rancho La Brea | M1 | no |
| UCMP | 215844 | Rancho La Brea | P2 | yes |
| UCMP | 215845 | Rancho La Brea | C | yes |
| UCMP | 215845 | Rancho La Brea | P2 | yes |
| UCMP | 215845 | Rancho La Brea | P3 | yes |
| UCMP | 215847 | Rancho La Brea | C | yes |
| UCMP | 215847 | Rancho La Brea | P2 | yes |

Larger and more robust than the modern wolf, the dire wolf was a highly successful predator that inhabited a wide range of habitats across the American continents during the Pleistocene (Dundas, 1999). Fossil evidence of the dire wolf spans several hundred thousand years, with the earliest record at least 252,000 years old (Mead et al., 1996). Although the terminal date of the dire wolf is not precisely known, its extinction at the end of the Pleistocene coincided with the abrupt extinction of numerous other megafaunal species in North America. The causes of this “Rancholabrean Termination” remain unclear, although major climate change and human predation/competition have been suggested (Haynes, 2008).

### Rancho La Brea tar pits

Fourteen teeth in the sample were obtained from the Rancho La Brea site collection at the University of California Museum of Paleontology (UCMP) in Berkeley, California. Rancho La Brea is located on the Santa Monica Plain in the northwest part of the Los Angeles Basin in southern coastal California, where heavy petroleum from a deep underground reservoir seeps to the earth’s surface along faults and permeable zones (Akersten et al., 1983:211). Over the last 40,000 years, these asphaltic sediments have accumulated and preserved thousands of Pleistocene megafaunal remains, representing one of the largest and most well-known fossil deposits in North America. Since 1906, excavations by the University of California and other institutions have recovered large quantities of these remains for scientific study, including thousands of dire wolves (Marcus, 1960).

The RLB teeth are darkly stained by varying degrees from petroleum impregnation in asphaltic sediments (Fig. 2). In appearance the tooth enamel ranges in color from light to dark brown. Most teeth were fractured either before or after death, allowing hydrocarbons from the depositional environment to permeate the inner tissues. Fracture and wear surfaces also reveal dark brown discoloration of the dentine and pulp. All teeth in the sample were removed *in situ* from mandibles, and only in cases where the tooth was loose enough to be pulled from its bony socket without damage. Bones associated with the teeth exhibit dark brown discoloration, as noted in previous studies (Cook et al., 1962; Ho et al., 1969) (Fig. 2).

**Figure 2.**
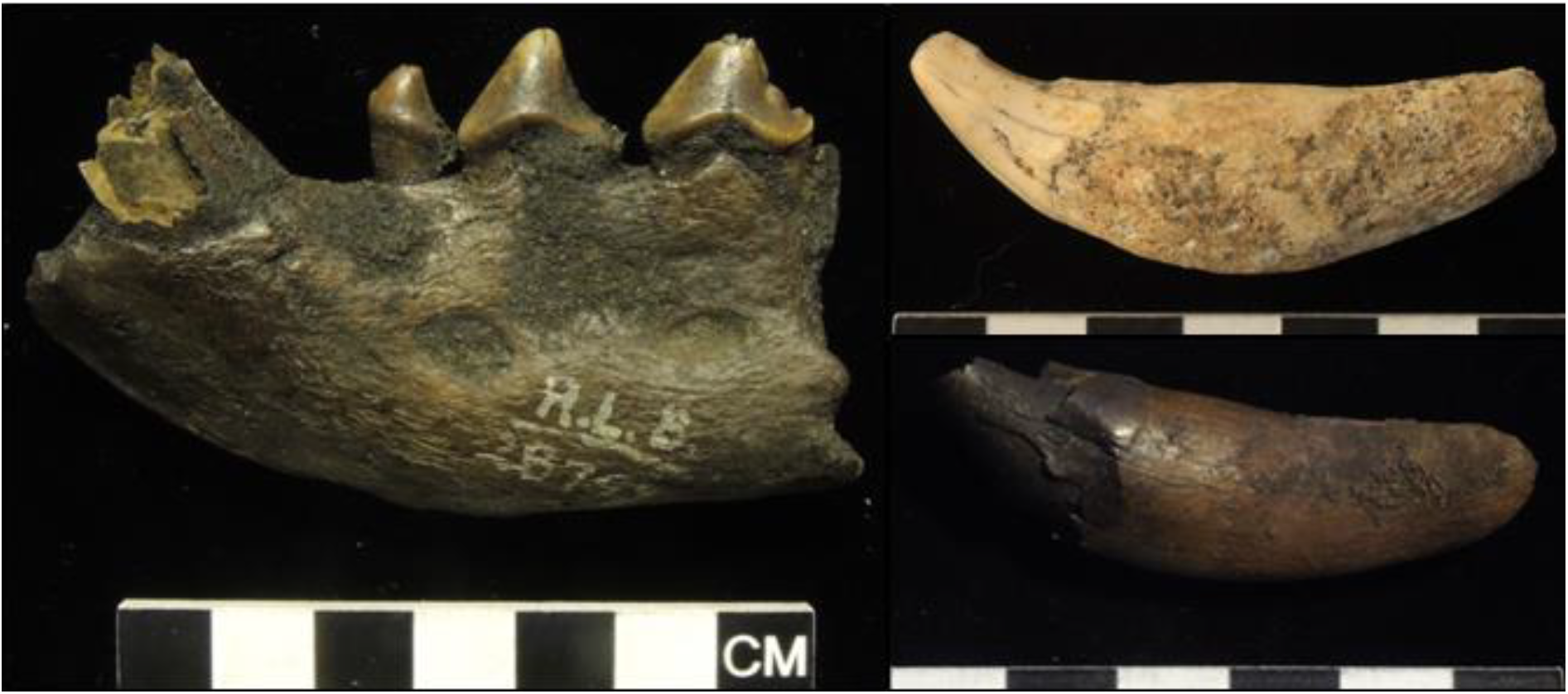
Photographs of teeth used in this study (scale bars are 2 cm). Fossils from the Rancho La Brea (RLB) tar pits display dark discoloration from petroleum impregnation (UCMP 215837, left; UCMP 215840, top right). Fossils deposited in the karstic sediments in the Cutler Hammock (CH) sinkhole exhibit little or no surface discoloration (FLMNH 276782, bottom right).

### Cutler Hammock sinkhole

Ten of the teeth in the sample were obtained from the Cutler Hammock site collection at the Florida Museum of Natural History (FLMNH) in Gainesville, Florida. Discovered in 1985, the Cutler Hammock site is a fossil deposit found in a large sinkhole in a Miami limestone formation, located on the Atlantic Coastal Ridge in southernmost peninsular Florida (Emslie and Morgan, 1995). Since its formation in the late Pleistocene (Rancholabrean Age), the Cutler Hammock sinkhole has filled with sediments and limestone rock. Amphibian remains from the site suggest that water was available in the sinkhole for at least part of the year during the formation of the deposits (Emslie and Morgan, 1995). With at least 42 individuals represented, this site contains the largest fossil assemblage of dire wolf remains in North America after Rancho La Brea (Morgan and Emslie, 2010).

On the outside, the CH teeth exhibit minimal effects from the karstic environment of the sinkhole (Fig 2). The tooth enamel appears mostly normal in color, ranging from yellowish to grayish white. Fracture surfaces on most teeth reveal a similar color range for the dentine, although in some cases the dentine is brown and/or poorly preserved. All of the selected CH teeth were isolated, and physically unassociated with any mandible or mandibular fragments present in the CH collection.

## 5. METHODS

### Sample preparation

For destructive analysis, teeth were embedded in a greased EPDM rubber mounting cup with a 5:1 resin:hardener mixture of Buehler epoxide, and cured overnight at room temperature. Within the hardened resin block, each tooth was then sectioned at a longitudinal plane through the highest point of dentine under the cuspal enamel using an IsoMet low speed saw with a diamond wafering blade. Thick sections (approx. 2-3 mm) were ground with abrasive discs of decreasing grit size (P80-P1200) and polished with diamond compound paste. After approximately 30 second of sonication in distilled water, section surfaces were etched with 2M HCl for 3-5 seconds.

### Scanning electron microscopy (SEM)

A TM3000 Tabletop SEM (Hitachi) operating at 15 kV were used for back-scatter mode imaging and energy-dispersive spectroscopy (EDS) analysis of a) tooth samples in sectioned blocks treated with carbon coating, and b) fractured surfaces of non-sectioned non-coated teeth.

### Light microscopy

For light microscopy, thin sections (30 μm) were prepared by the Thin Section Lab in Toul, France. Both thin and thick sections (etched and non-etched with polishing) were viewed and photographed with and without crossed polarizing filters using a Nikon light microscope equipped with a Leica digital camera.

### X-ray diffraction

Minute scrapings of enamel from five teeth (modern gray wolf, La Brea samples UCMP 215834 and15845, and CH samples FLMNH 276788 and 276780) were put on a glass spindle, and X-ray diffractograms were recorded using a Rigaku R-Axis Spider X-ray unit operating with a molybdenum anode at 50kV/40mA. Data was recorded for five minutes per sample and analyzed using the JADE v8.2 software.

## 6. RESULTS

Non-distorted optical microscope and SEM images were obtained for dire wolf teeth from the RLB and CH sites. The enamel microstructure of the RLB and CH teeth did not exhibit any clear differential effects of taphonomy. In both samples, prisms, prism rods, and Retzius lines could be seen both with SEM and light microscopy (Fig. 3). Hunter-Schreger bands were clearly visible in all teeth, although for one CH tooth (FLMNH 276797), the bands were not visible under normal (non-polarized) light (Fig. 4). In both RLB and CH enamel, cross-striations were visible in thin sections under cross-polarized light, as well as in fractured surfaces with SEM (Fig. 5). No cross-striations were observed in the polished cross-section samples, neither with optical microscopy nor with SEM.

**Figure 3.**
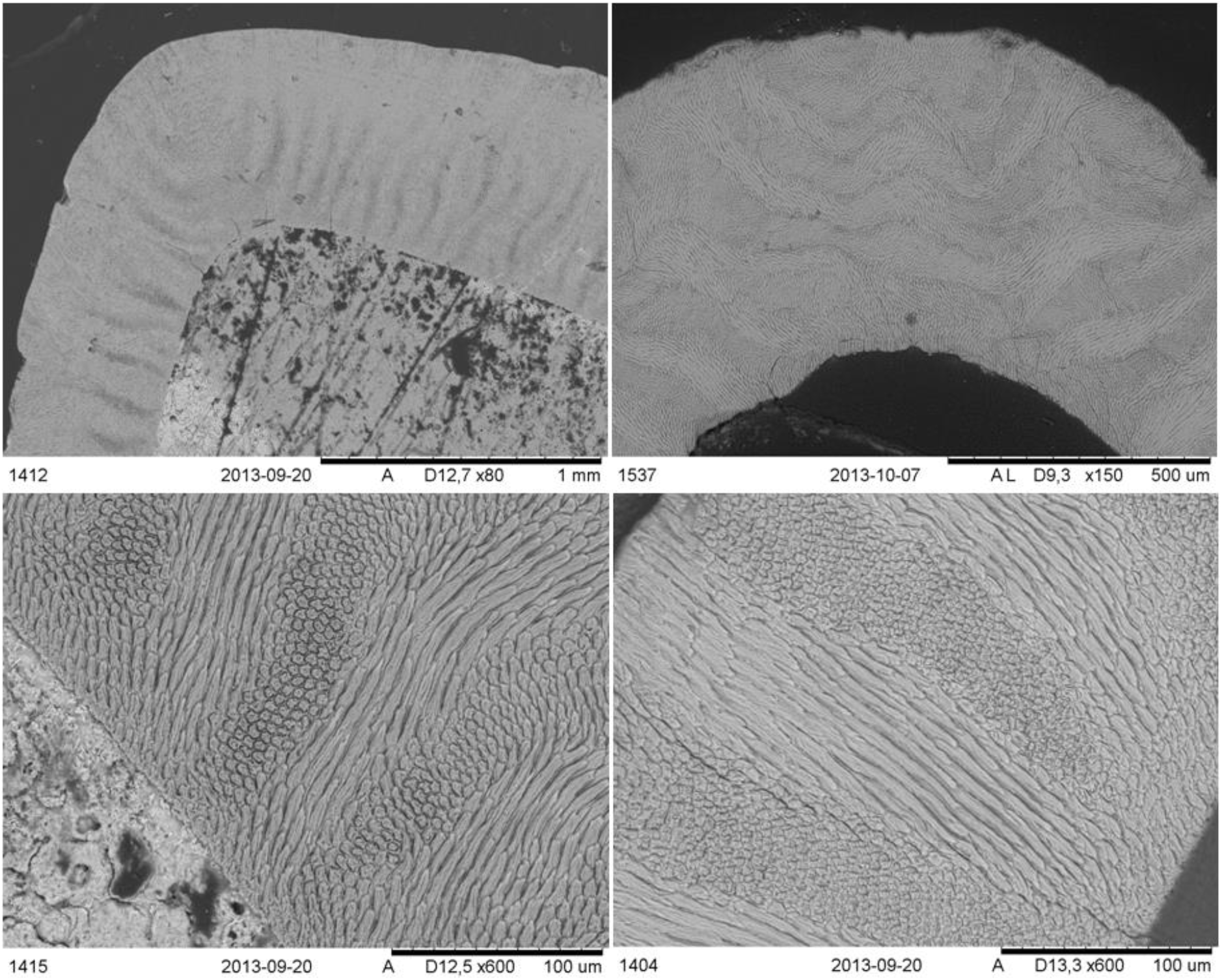
SEM backscatter images of teeth from the Cutler Hammock (CH) sinkhole (FLMNH 276788, top and bottom left) and the Rancho La Brea (RLB) tar pits (UCMP 215842, top and bottom right), readily allowing observation of Hunter-Shreger bands at the tip (top row) and sides (bottom row) of the tooth crown.

**Figure 4.**
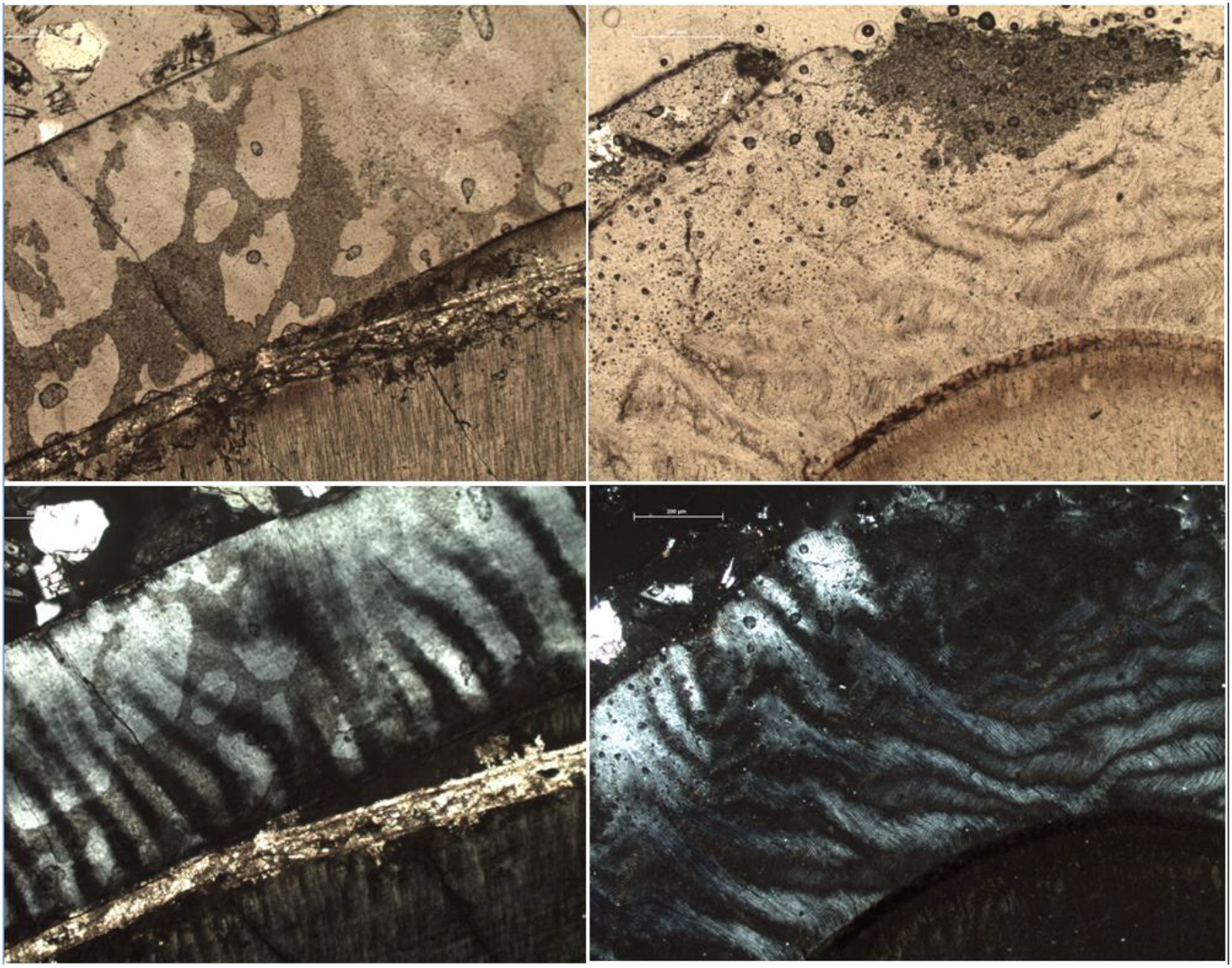
Optical microscope images of teeth from the Cutler Hammock (CH) sinkhole (FLMNH 276797, top left and bottom left) and the Rancho La Brea (RLB) tar pits (UCMP 215834, top right and bottom right). Under non-polarized light (top row) exogenous staining in the enamel of the Cutler Hammock tooth is clearly visible. Under fully cross-polarized light (bottom), this staining interferes with identification of Hunter-Schreger bands in the CH tooth.

**Figure 5.**
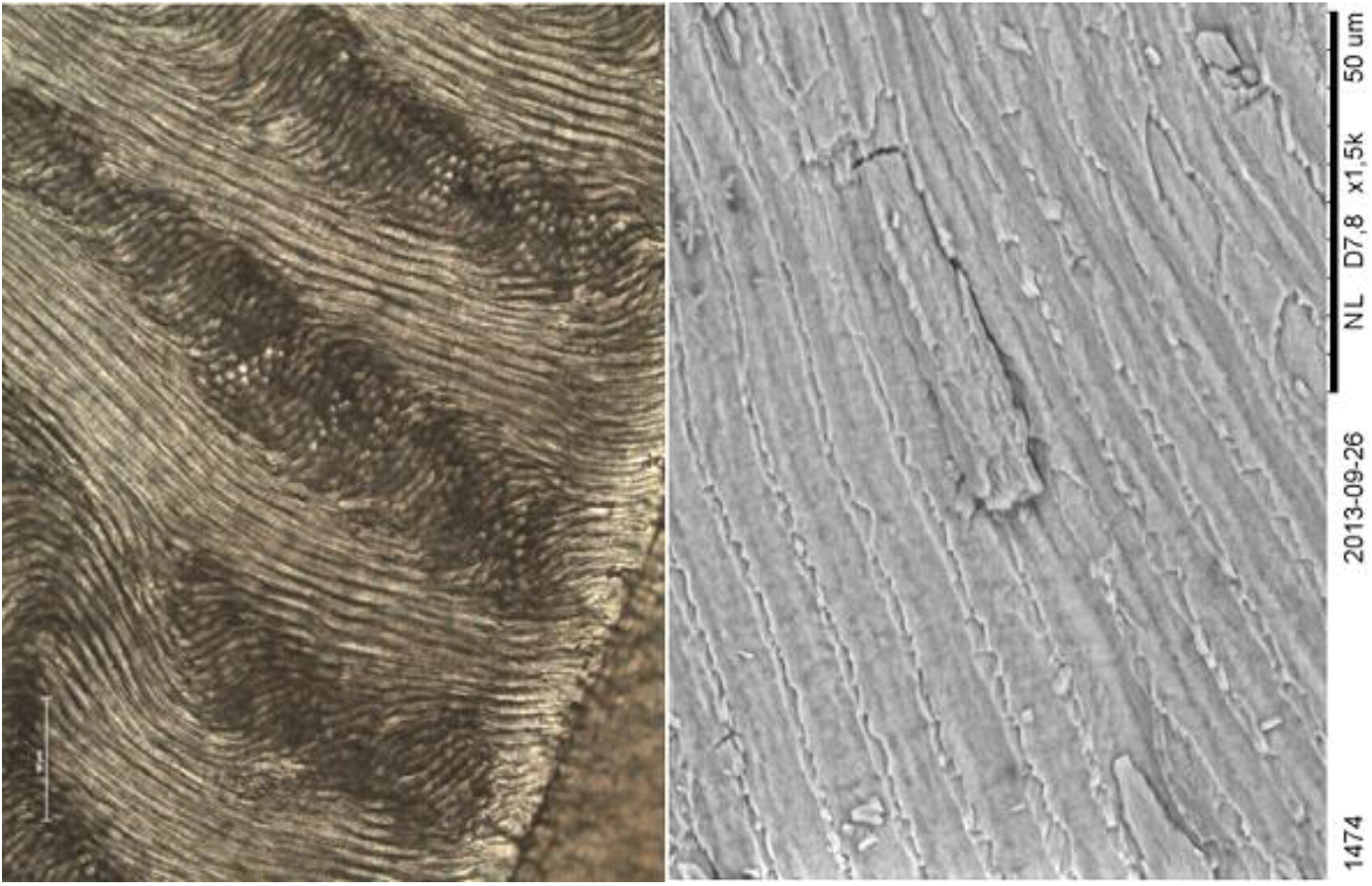
Enamel prism cross striations in teeth from the Rancho La Brea (RLB) tar pits, observed with non-polarized transmitted light on a thin section (UCMP 215834, left) and with SEM on a non-coated fractured surface (UCMP 215242, right).

For one of the CH teeth, SEM imaging revealed enamel areas with lower mineral density. Elemental analysis (SEM-EDS) revealed increased carbon and decreased phosphorus levels in these regions, indicating that the tooth apatite had been replaced with calcite. This replacement does not appear to affect the enamel microstructure. No alterations of chemical composition were observed in the enamel from the RLB samples. These results prompted us to use XRD analysis to investigate the crystallinity of the enamel apatite in two RLB, two CH, and one modern wolf tooth. From Fig. 6 it is clear that although all diffractograms show peaks corresponding to apatite, the signal intensities are strongest for the modern wolf tooth and the teeth from Rancho La Brea. In contrast, the two CH enamel samples display XRD peaks that are not only weaker but also broader, indicating compromised crystallinity. Earlier studies have demonstrated that the combination of SEM-EDS and XRD analysis is a very powerful approach for detailed characterization of inorganic materials (Roos et al., 2020; Scott et al., 2009; Smith et al., 2015). Here, the combined XRD results (reduced crystallinity) and SEM-EDS results (altered elemental composition) clearly show that the enamel in the CH samples has lost some of its chemical integrity.

**Figure 6.**
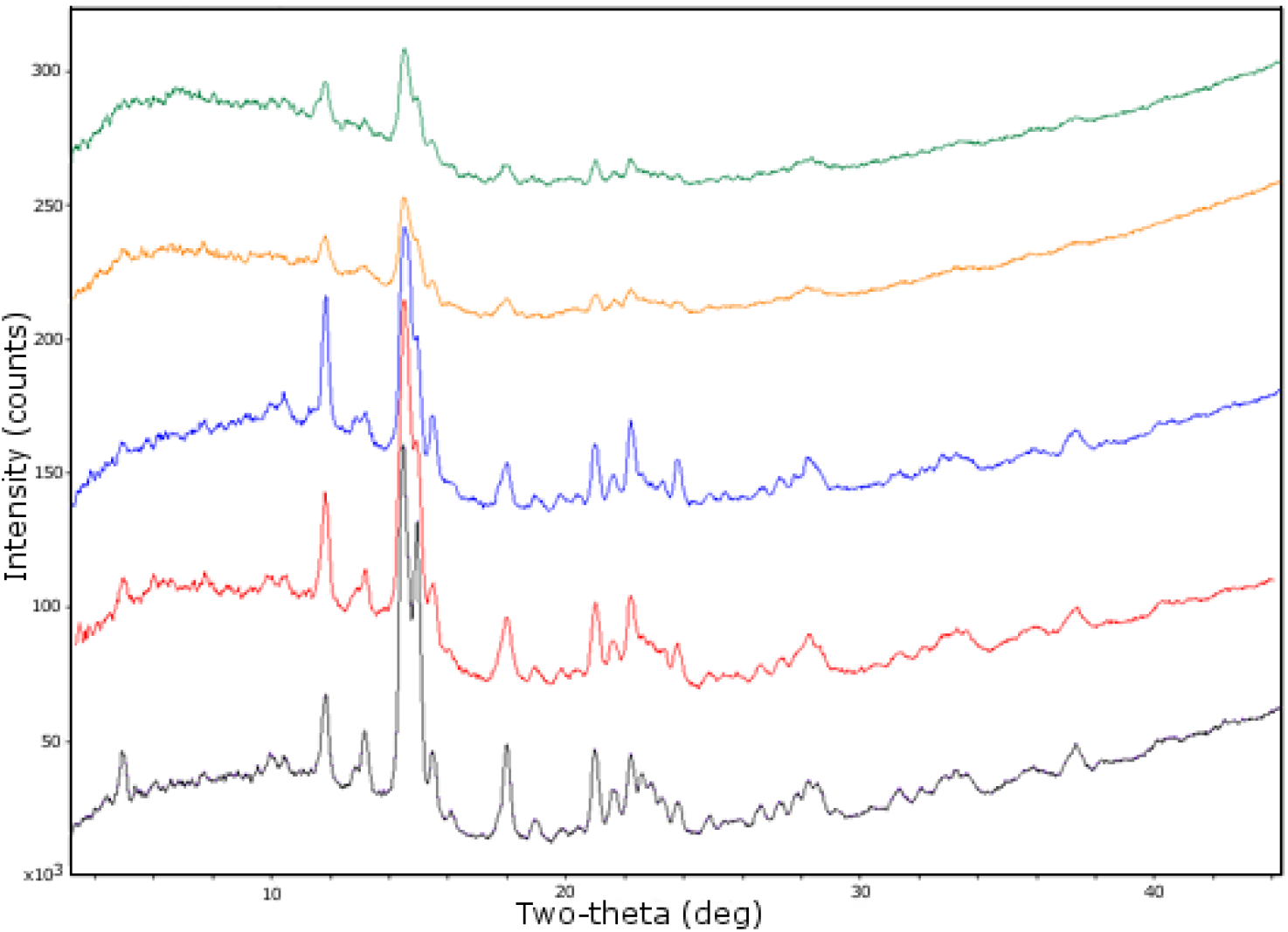
XRD patterns for samples of tooth enamel from: a modern wolf (black), two RLB dire wolf samples (UCMP 215834, red; UCMP 215845, blue), and two CH dire wolf samples (FLMNH 276788, orange; FLMNH 276780, green). The spectra are shown without baseline correction, and the similar baseline distortion (originating from the glass spindle) shows the similarity in sample preparation and data recording.

Dentine preservation was very different between the CH and RLB samples (Fig. 7–8). The RLB dentine was consistently brown in color, but microstructures such as Andersen’s lines and dentinal tubules were clearly observable with both SEM and optical microscopy. In all the CH teeth the analyzed dentine varied in color, from bright orange to white, and contamination and/or significant alteration was evident at the tissue and microscopic levels. Microstructural features such as Andersen’s lines and dentinal tubules were well preserved in one sectioned CH tooth (FLMNH 276797), but were not consistently observed in the others. Differences in dentine between the two samples were also shown by EDS, which detected the presence of 1-3 wt.% iron in one CH tooth (FLMNH 276788) (Fig. 9) and an elevated level of fluorine, 1.9 wt.%, in another (FLMNH 276780). The iron in the CH tooth is most likely iron oxide in the form of haematite, Fe2O3, as indicated by the reddish-orange appearance of the dentine region (Fig. 7). The RLB dentine did not contain any elements inconsistent with apatite apart from small amounts (0.69-0.96 wt.%) sulfur, which is found in both petroleum and tooth collagen.

**Figure 7.**
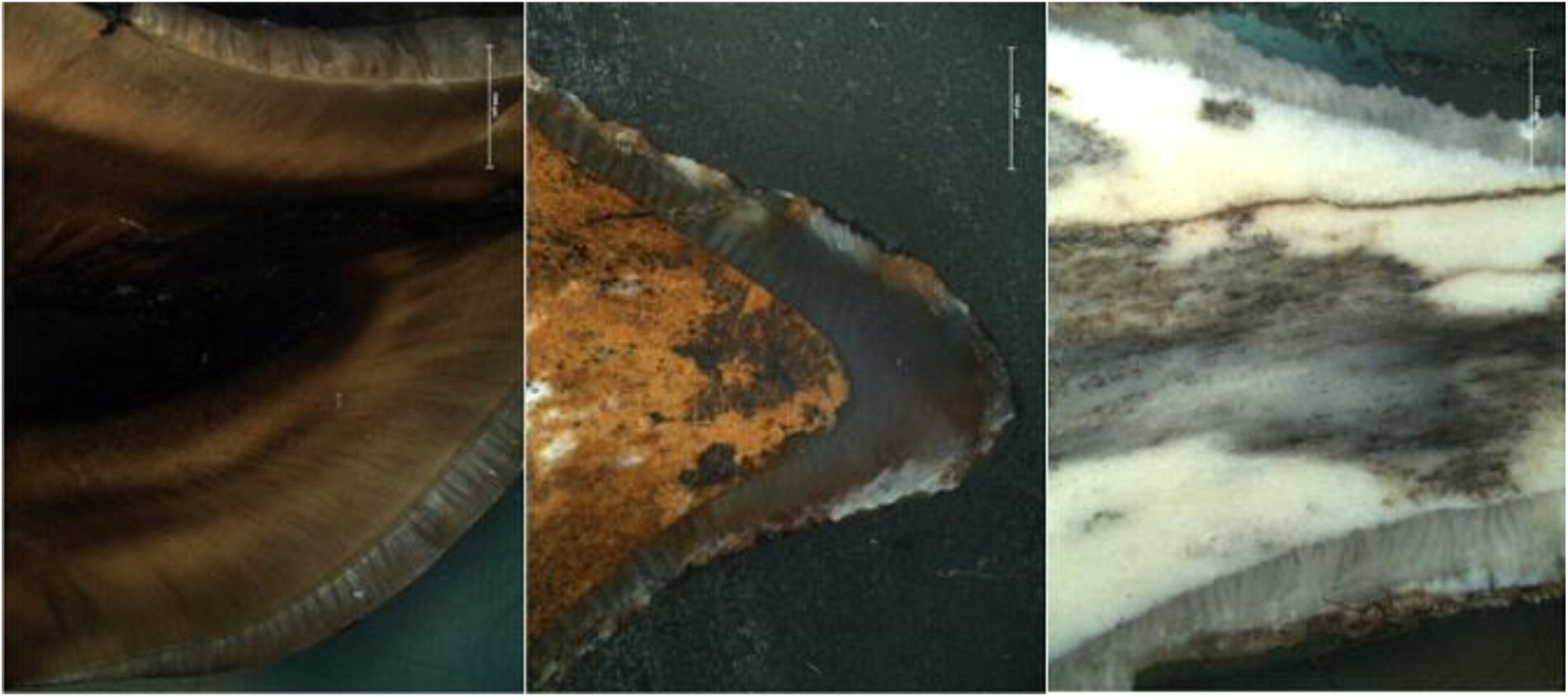
Optical microscope images of polished cross-sections of teeth from the Rancho La Brea (RLB) tar pits (UCMP 215845, left) and the Cutler Hammock (CH) sinkhole site (FLMNH 276788, middle; FLMNH 276780, right), recorded under reflected light.

**Figure 8.**
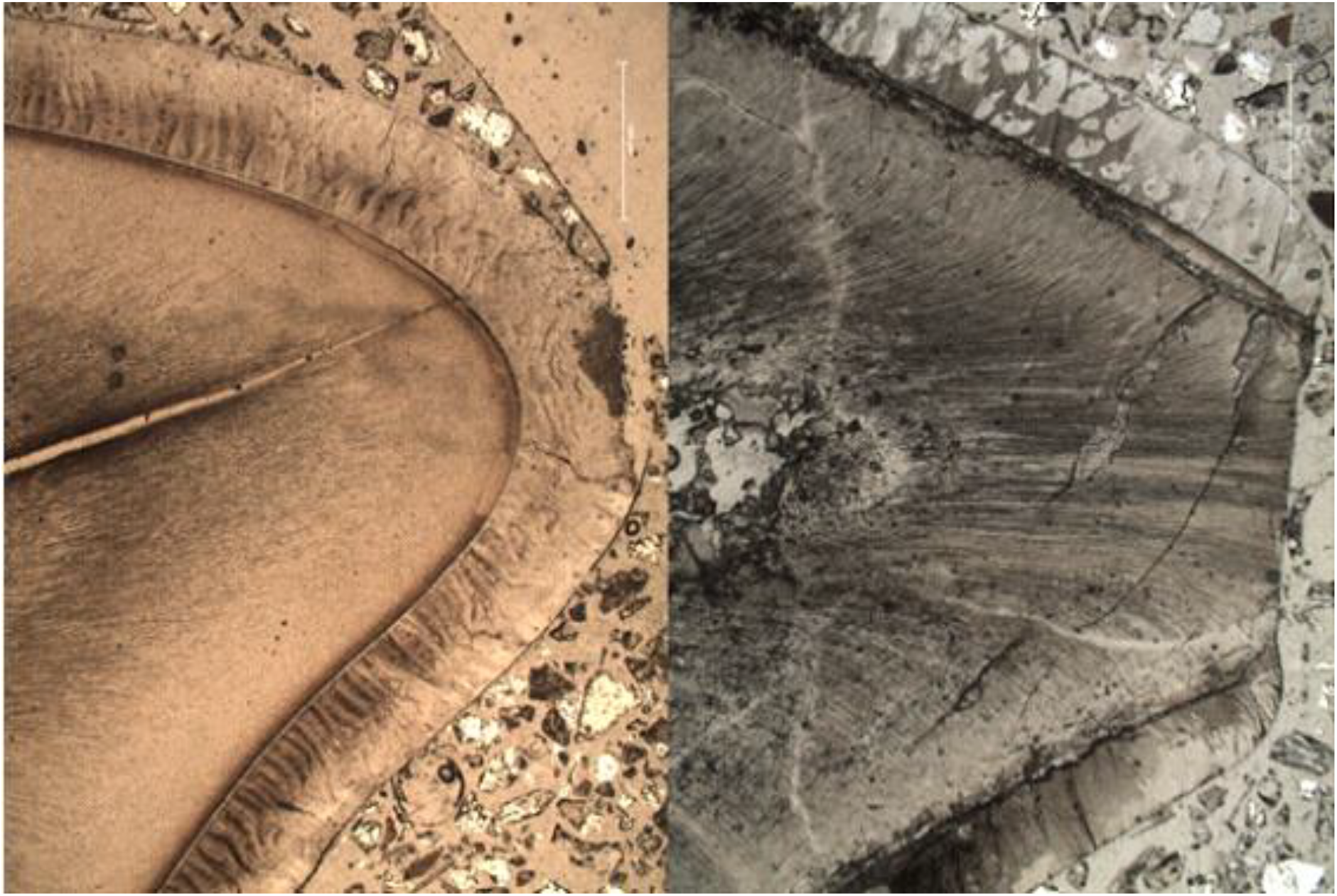
Optical microscope images of thin sections of teeth from the Rancho La Brea (RLB) tar pits (UCMP 215834, left) and the Cutler Hammock (CH) sinkhole (276797, right), recorded under non-polarized transmitted light. In the Cutler Hammock tooth, significant taphonomic alterations of the dentine can be seen in comparison to the Rancho La Brea tooth.

**Figure 9.**
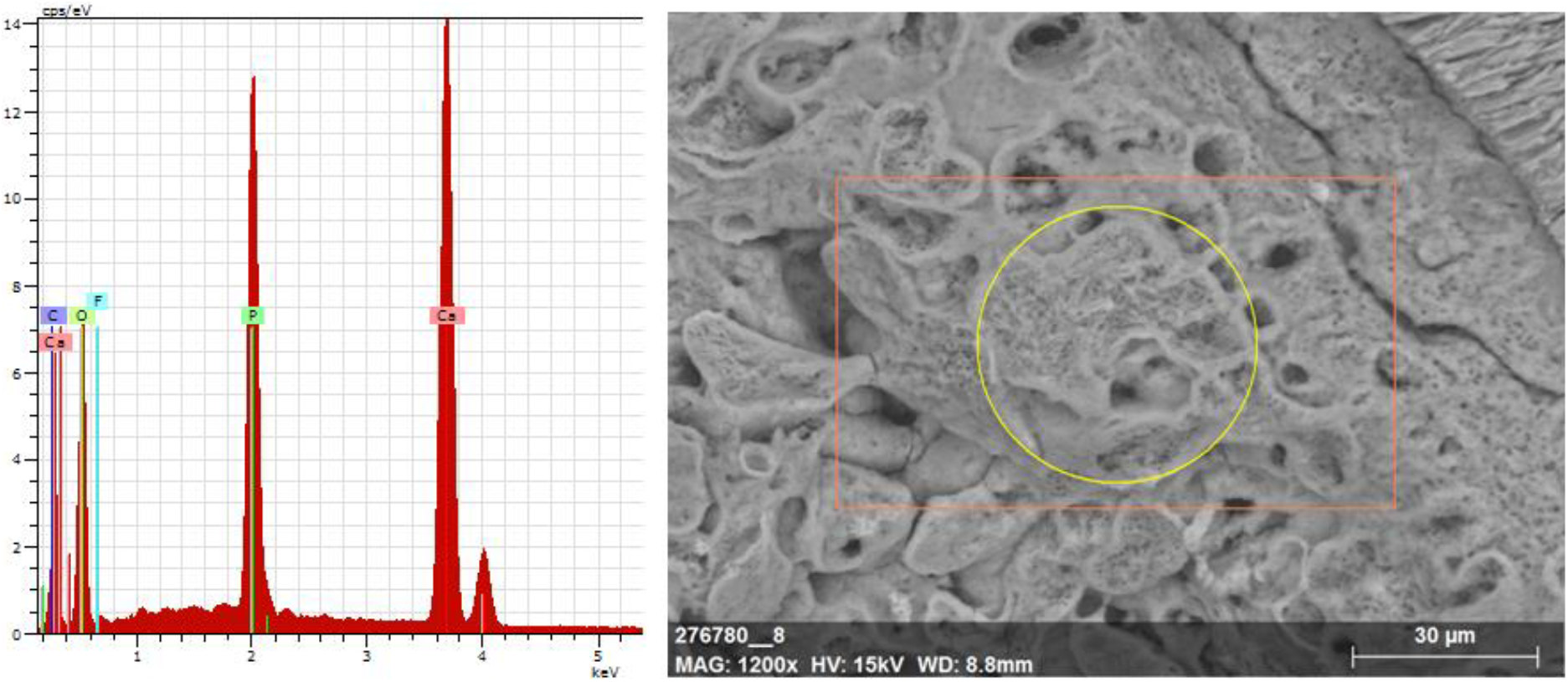
SEM-EDS spectrum and associated SEM image of a Cutler Hammock (CH) tooth cross-section (FLMNH 276788), showing elevated levels of iron in the dentine.

## 7. DISCUSSION

By visual inspection, it is immediately clear that fossils from Rancho La Brea are stained by the black petroleum. Cross-sections showed this dark staining to permeate the entire teeth, as the dentine in the RLB teeth were altered from their natural yellow-orange color to a dark brown hue. Yet, all microscopic structures could be readily observed both in the dentine and enamel, indicating an absence of taphonomic effects in the RLB teeth except for optical discoloration. Not even the polarizing properties of the enamel crystals were affected by the discoloration, as incremental markings were readily visible in light microscopy.

For the teeth from the Cutler Hammock site, cross-sections revealed staining in various colors as the dentine ranged from strong orange-red to an unnatural bright white. These discolorations were consistent with, and arguably explained by, the elemental analysis that revealed increased levels of e.g. iron and fluorine in the CH teeth. Such alterations are in line with previous analyses of fossils from karstic deposits (Michel et al., 1996), suggesting that they are related to the geochemical conditions in the Cutler Hammock sinkhole. As the higher porosity of the dentinal collagen-apatite matrix provides more open pathways for aqueous intrusion, fluids within a burial environment have been shown to permeate this tissue more efficiently than enamel (Dauphin and Williams, 2007). Taphonomic effects on teeth can therefore result in high variability in dentine preservation, possibly with little remaining dentine structure and opacity in light microscope cross sections (Hillson, 2005:190). Accordingly, other detected elements such as iron most likely also contaminated the dentine via fluvial infiltration through the pores.

These findings may not appear very surprising, as enamel generally is the most stable part of the tooth due to a low content of organic matrix (Dauphin and Williams, 2007). However, spectroscopic analysis revealed a loss of chemical integrity for the CH tooth enamel, related to partial replacement of tooth apatite with calcite. While this replacement appears to preserve the enamel microstructure, in line with established principles for mineral fossilization, it suggests that neither dentine nor enamel of CH teeth is suitable for chemical analysis, even though their outward appearance showed no abnormalities in color or condition. In contrast, the RLB teeth showed minimal alteration despite the petroleum staining, making them potentially suitable for both histological and chemical analysis. Thus, the interior integrity of the studied teeth is clearly not correlated with their exterior appearance.

The obtained results can arguably be rationalized in terms of petroleum chemistry. Being both viscuous and hydrophobic/lipophilic, petroleum is very sticky, and its hydrocarbons readily adhere to bone and tooth tissues. Due to their low reactivity, these hydrocarbons do not interact chemically with the collagen or apatite material in tooth/bone. Instead, by permeating the teeth the petroleum plugs their pores, preventing the kind of fluvial processes that caused chemical replacement in the CH sample. Fluvial processes are further prevented through the general hydrophobicity of hydrocarbons, which keeps the petroleum seeps relatively water-free and also anaerobic. Both organic and inorganic skeletal material encased in such seeps is therefore well protected against diagenetic effects and likely readily accessible for analysis of biomolecules such as proteins and deoxyribonucleic acids (DNA), in line with previous reports that petroleum impregnation is an ideal preservative for bone collagen (Ho et al., 1969) that does not effect molecular or structural change in organic material (Shelton, 1994). The presence of petroleum will however bias any study that involves measuring isotope ratios of the elements typically found in hydrocarbons, most notably carbon but to a lesser extent also oxygen and nitrogen. For example, being a material of infinite geological age petroleum lacks ^14^C isotopes, wherefore petroleum-impregnated bones and teeth cannot be radiocarbon dated without first separating and removing the petroleum (Ho et al., 1969; Jull et al., 2004). On the other hand, fossils from petroleum seeps should be well suited for analysis of isotope and trace elements profiles of heavier elements such as the transition metals.

In conclusion, we here demonstrate that fossil dire wolf teeth from a petroleum seep exhibit consistently well-preserved dental tissues, despite being discolored. In contrast, dire wolf teeth of similar age from a karstic sinkhole displayed a flawless outward appearance but poor chemical preservation, as the dental tissues had been compromised by exogeneous elements via fluvial intrusion. Thus, petroleum seeps appear to provide a suitable environment for preserving the chemical integrity of fossil biological material.

## ACKNOWLEDGMENTS

The authors wish to thank Pat Holoroyd (UCMP) and Richard Hulbert (FLMNH) and other museum personnel for assistance and permission to access the samples, Kjell Jansson and Dan Zetterberg (Stockholm University) for access to microscope facilities, and the Smithsonian’s Dept. of Mineral Sciences for help with the chemical analysis.

